# Rates of increase of antibiotic resistance and ambient temperature in Europe: a cross-national analysis of 28 countries between 2000-2016

**DOI:** 10.1101/414920

**Authors:** Sarah F. McGough, Derek R. MacFadden, Mohammad W. Hattab, Kåre Mølbak, Mauricio Santillana

## Abstract

**Background:** Widely recognized as a major public health threat globally, the rapid increase of antibiotic resistance in bacteria could soon render our most effective method to combat infections obsolete. Factors influencing the burden of resistance in human populations remain poorly described, though temperature is known to play an important role in mechanisms at the bacterial level.

**Methods:** We performed an ecologic analysis of country level antibiotic resistance prevalence in 3 common bacterial pathogens across 28 countries in Europe, and used multivariable models to evaluate associations with minimum temperature and other predictors over a 17-year period (2000-2016). We quantified the effects of minimum temperature, population density, and antibiotic consumption on the rate of change of antibiotic resistance across geographies.

**Findings:** For three common bacterial pathogens and four classes of antibiotics, we found evidence of a long-term effect of ambient minimum temperature on rates of increase of antibiotic resistance across 28 countries in Europe between 2000-2016. Specifically, we show that across all antibiotic classes for the pathogens *E. coli* and *K. pneumoniae*, European countries with 10°C warmer ambient temperatures have experienced more rapid increases in antibiotic resistance over the 17-year period, ranging between 0.33%/year (95% CI 0.2, 0.5) and 1.2%/year (0.4, 1.9), even after accounting for recognized drivers of resistance including antibiotic consumption and population density. We found a decreasing relationship for *S. aureus* and methicillin of −0.4%/year (95% CI −0.7, 0.0), reflecting widespread declines in MRSA across Europe over the study period.

**Interpretation:** Ambient temperature may be an important modulator of the rate of change of antibiotic resistance. Our findings suggest that rising temperatures globally may hasten the spread of resistance and complicate efforts to mitigate it.

**Funding:** Canadian Institutes of Health Research Fellowship (D.R.M.).

## Research in context

### Evidence before this study

While multiple studies have independently pointed out associations between temperature and bacterial biological cycles, few studies have focused on characterizing the relationship of ambient temperature and antibiotic resistance on a multi-geography or multi-country scale. We performed a general search on MEDLINE using the joint keywords “antibiotic resistance” and “climate” for original research published up to June 22, 2018. A recent study conducted in the US identified that locations with warmer ambient temperatures experienced higher values of antibiotic resistance prevalence using three years of data, and one European country-level analysis discovered that higher distance from the equator was a predictor for lower risk of *Acinetobacter spp.* resistance to carbapenems.

### Added value of this study

For the first time, we present evidence that ambient temperatures may modulate the rate of increase of antibiotic resistance across Europe. Using a comprehensive dataset containing information across 28 countries, for 17 years, 3 bacterial pathogens, and 4 antibiotic classes collectively representing over 4 million tested isolates, we show that antibiotic resistance has increased more rapidly in warmer regions over a period of nearly 2 decades. This finding helps us better understand the currently observed geographic patterns of antibiotic resistance prevalence in Europe that show that warmer countries experience higher prevalence of antibiotic resistance, and may explain what has been recently observed in the US.

### Implications of all the available evidence

Our findings motivate the need for further studies to better characterize the role of temperature as a mechanistic modulator of transmission of antibiotic resistant microbes, either at the microscopic level or at the population level via human activities. With ambient temperatures rising in our planet, our results suggest that we may enter a post-antibiotic era sooner than current estimates suggest.

## Introduction

For almost a century, antibiotics have been our most effective way of combating bacterial infections, and underpin enormous population health gains. However, soon after their initial introduction, bacterial pathogens demonstrated their propensity to acquire and propagate mechanisms to withstand the effects of these agents. Decades of unfettered use of antibiotics has been recognized as a main driver behind the selection and spread of resistant bacteria globally.^1,2^

Antibiotic resistance is one of the world’s greatest public health threats today, with the potential to render many existing classes of antibiotics ineffective in the near future.^3,4^ To combat this crisis, numerous national and international bodies have begun to develop policy and fund research aimed at identifying and targeting drivers of resistance.^3,5^ However, we have an incomplete understanding of the factors beyond antibiotic consumption that influence the distribution and spread of antibiotic resistance in human populations.

Temperature is one of the strongest drivers of bacterial reproduction and can also modulate aspects of horizontal gene transfer through which resistance genes can be exchanged.^6,7^ On a population level, ambient (air) temperature has been associated with rates of human carriage,^8^ and influences the intensity of human activities, such as food animal production, that could promote increased use of antibiotic agents. Thus, directly or indirectly, temperature has the potential to modify the process of bacterial transmission, transfer of mobile elements between bacteria, and selection of antibiotic resistant organisms at bacterial and human population scales. The effects of warming temperatures on a variety of infectious diseases globally have been identified by the World Health Organization; however, the impact of climate change on the distribution of antimicrobial resistance has been relatively ignored.^9^ A recent ecologic study evaluated the distribution of antibiotic resistance in common bacterial pathogens across the United States during the years 2013-2015, and found that antibiotic resistance prevalence was linked to local minimum ambient temperatures across geographies.^10^ However, due to limited availability of historical data, the study could not demonstrate the temporal effects of climate on antibiotic resistance.^10^

Here we use one of the most comprehensive antibiotic resistance databases in existence^11,12^ to identify if the rates of change of antibiotic resistance across countries in Europe, over the years 2000-2016, may have been modulated by ambient temperature, and whether these findings may explain observed associations between temperature and antibiotic resistance across geographies.

## Methods

### Study Design

We performed an ecologic analysis of the time evolution of country-level antibiotic resistance prevalence in common bacterial pathogens across 28 countries in Europe, and evaluated associations with temperature and other predictors over a 17-year period (2000-2016). We built multivariable linear models of antibiotic resistance prevalence, for 3 common pathogens and up to 4 different classes of antibiotics. The study assessed 28 European countries: Austria, Belgium, Bulgaria, Croatia, Czech Republic, Denmark, Estonia, Finland, France, Germany, Greece, Hungary, Iceland, Ireland, Italy, Latvia, Lithuania, Luxembourg, Netherlands, Norway, Poland, Portugal, Romania, Slovakia, Slovenia, Spain, Sweden, and the United Kingdom.

### Antibiotic resistance data

Antibiotic resistance data at the country level were collected as part of national surveillance for 28 countries across Europe, for three common bacterial Gram-positive and Gram-negative pathogens: *Escherichia coli, Klebsiella pneumoniae*, and *Staphylococcus aureus*, representing approximately 3.4 million, 0.54 million, 0.52 million tested isolates, respectively. Antibiotic resistance was identified as prevalence of resistance (% resistant) among reported isolates for a given year, country, and antibiotic class. Antibiotic susceptibility was evaluated, where available, for the following classes: aminopenicillins (*E. coli*), 3rd generation cephalosporins (*E. coli* and *K. pneumoniae*), fluoroquinolones (*E. coli* and *K. pneumoniae*), aminoglycosides (*E. coli* and *K. pneumoniae*), and methicillin (*S. aureus*). Antibiotic resistance data originate from the European Antimicrobial Resistance Surveillance Network (EARS-NET).^12^ This publicly available database includes national annual human antibiotic resistance data across common bacterial pathogens for common drugs, for the majority of European countries over time periods dating back to 2000. These national data comprise one of the most comprehensive antibiotic resistance datasets in existence though constrain the analysis to the country level.

### Predictors and confounders

*Antibiotic consumption* here refers to either sales data, reimbursement data, or both, depending upon the country. We do not distinguish between these two sources, rather referring to them both as ‘consumption’. To account for country level differences in antibiotic consumption, we used annual country level antibiotic consumption data from the European Centers for Disease Control (ECDC), specifically prescribing rates expressed as the number of defined daily doses (DDD) per 1000 inhabitants per day, from combined inpatient and outpatient sources (where available) for all major Anatomical Therapeutic Chemical Classification System antibiotic classes. These data from The European Surveillance System – TESSy, were provided by Austria, Belgium, Bulgaria, Croatia, Czech Republic, Denmark, Estonia, Finland, France, Germany, Greece, Hungary, Iceland, Ireland, Italy, Latvia, Lithuania, Luxembourg, Netherlands, Norway, Poland, Portugal, Romania, Slovakia, Slovenia, Spain, Sweden, and the United Kingdom, and released by the ECDC. For each country, these data were available for up to 17 years, from 2000 to 2016. Antibiotic prescribing data were missing for some countries and years, and coverage increased over time. The following antibiotics were represented by the following ATC codes: Penicillins - J01C, Fluoroquinolones - J01M, Aminoglycosides - J01G, and 3rd Generation Cephalosporins - J01DD.

Our second predictor was ambient temperature. Given that *minimum temperature* has been identified as important when describing species survival^13,14^ and was found to have meaningful associations with antibiotic resistance prevalence in a previous ecological study,^10^ we focused on extracting this attribute. Constrained by the spatial resolution of the antibiotic resistance data to a country-level analysis and noting that certain European countries possess large north-south latitudinal ranges or topographical variations that may limit human population settlements, we used three different approaches to calculate annual minimum temperature values for our study. First, we calculated the country-level annual average minimum temperature using modeled and assimilated meteorological data, available at a native geographic resolution of 0.5° × 0.625° from the Modern-Era Retrospective analysis for Research and Applications, Version 2 (MERRA-2).^15^ The MERRA-2 data are publicly available through the Global Modeling and Assimilation Office (GMAO) at NASA Goddard Space Flight Center. MERRA-2 contains hourly information from 1980 through the present date, with no missing data and geographic coverage. This consistency over time and space makes MERRA-2 an ideal data source for this analysis. For this approach, we extracted the daily minimum temperature for the 17-year period (2000-2016) from MERRA-2 gridded rasters onto a European shapefile containing country-level attributes, took the mean of daily minimum temperature across grid cells spanning each country, and computed the annual mean, for each country, as the mean of daily mean minimum temperature over the calendar year. The shapefile was obtained from the Eurostat Geographic Information System of the Commission (GISCO) of the European Commission, and geodata extractions were performed with the aid of the “rgdal” and “raster” packages in R (Version 3.3.2).

The other two approaches to calculate minimum temperatures were conceived to assess the sensitivity of our findings to our choice of climate data source and spatial resolution. For these two subsequent approaches, we used city-level temperature data obtained from (a) MERRA-2 data and (b) local weather stations. For (a), daily minimum temperature from MERRA-2 were obtained from the single 0.5° x 0.625° grid cell covering the centroid of each capital city, and the annual mean for each country was computed over the calendar year. For (b), we obtained annual average minimum temperature from the most complete weather station data corresponding to a capital or populous city, as provided by the European Climate Data & Assessment (ECD&A) project of the Royal Netherlands Meteorological Institute (KNMI). 26 of the 28 countries had available weather station data, and the list of cities used for each country can be found in Table S1. It is important to note that using weather station data removes the theoretical modeling assumptions implicit in reanalysis data such as MERRA-2.

We computed the population density (persons/km^2^) for each country and year using annual population estimates obtained through Eurostat. We included country as a predictor in order to capture country-level confounding effects.

### Data analysis

We initially assessed the pairwise associations between relevant predictors, including minimum temperature, population density, and antibiotic consumption, and antibiotic resistance across countries, pathogens, and antibiotic classes. Additionally, we plotted the relationship between minimum temperature and antibiotic resistance within levels of predictors (antibiotic consumption-median, and population density-tertile) to assess for potential confounding and effect modifications. Log scales were used for antibiotic consumption given the large spread of values. To ease visualization of the trend of antibiotic resistance across multiple different antibiotics, we centered the data about the mean and normalized by the standard deviation, by pathogen, antibiotic, and year. To further visualize temporal trends, we computed the slope for the effect, by year, of minimum temperature on antibiotic resistance as well as the rate of change of antibiotic resistance over the 17-year period. These were done for each pathogen type and class of antibiotic.

To visualize the distribution of antibiotic resistance and minimum temperature across Europe, we generated choropleth maps of 1) normalized antibiotic resistance across the 3 pathogens and 2) annual average minimum temperature, each summarized over the 17-year time series. To produce a summary measure of antibiotic resistance, we first computed the weighted average of antibiotic resistance for each year, normalized by pathogen and antibiotic class and weighted by the number of isolates tested for a given pathogen and antibiotic class. The simple average of these values across years was then used to visualize the average normalized antibiotic resistance per country between 2000-2016. We also mapped the annual average minimum temperature by country as the mean of annual minimum temperature over the 17 years. Maps and figures were generated using the “ggplot2” package in R.

### Statistical analysis

Multivariable linear models were used to measure the association between minimum temperature (°C) and antibiotic resistance prevalence, adjusting for country, year, population density (persons/km^2^), antibiotic consumption (prescriptions per 1000 inhabitants per day), and the interaction between time and minimum temperature. These models were replicated across each pathogen and antibiotic class, and had the structure:

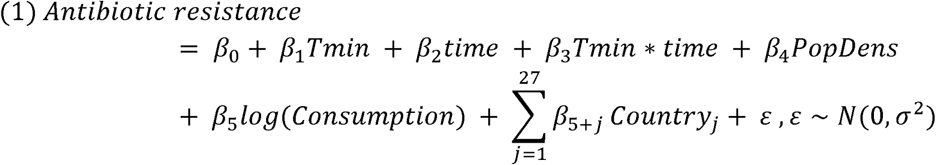

The interaction term between time and minimum temperature was included to assess the extent to which temperature is associated with the change in antibiotic resistance over time. To quantify the association of the temporal changes of antibiotic resistance prevalence with minimum temperature, we calculated the time derivative of Equation 1:

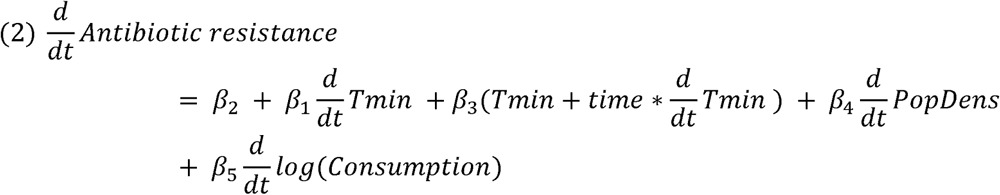

Equation 2 captures how the rate of change of antibiotic resistance evolves in time for each pathogen and antibiotic class. To determine the contribution of each of the terms to rates of change of antibiotic resistance, we estimated their values by approximating the time derivatives (of *Tmin*, *PopDens*, and *Consumption*) by the observed changes from year to year in the data. We then calculated the distributions of values for each individual term across countries and years. Given that *β* _2_ is constant across years and countries, we performed this step on the four rightmost terms of Equation 2.

## Results

Antibiotic resistance in *E.coli* and *K. pneumoniae* has increased over time for most countries in Europe (Figs. S1, S2), while *S. aureus* antibiotic resistance to methicillin has generally decreased over time (Fig. S3). Country-specific temporal trends in minimum temperature and antibiotic consumption show no clear tendencies as shown in Figs. S4 and S5. Countries have generally experienced steady increases or decreases in population density (Fig. S6).

For antibiotic resistance across all countries, years, pathogens, and antibiotic classes, we found evidence of a positive linear association with minimum temperature (Fig. 1), and observed that the relationship between temperature and resistance over geographies strengthened over time (Figs. 2A-C; Figs. S7A-C; Figs. S8A-C). Such evidence is consistent with previous findings across the US^10^.

**Figure 1:**
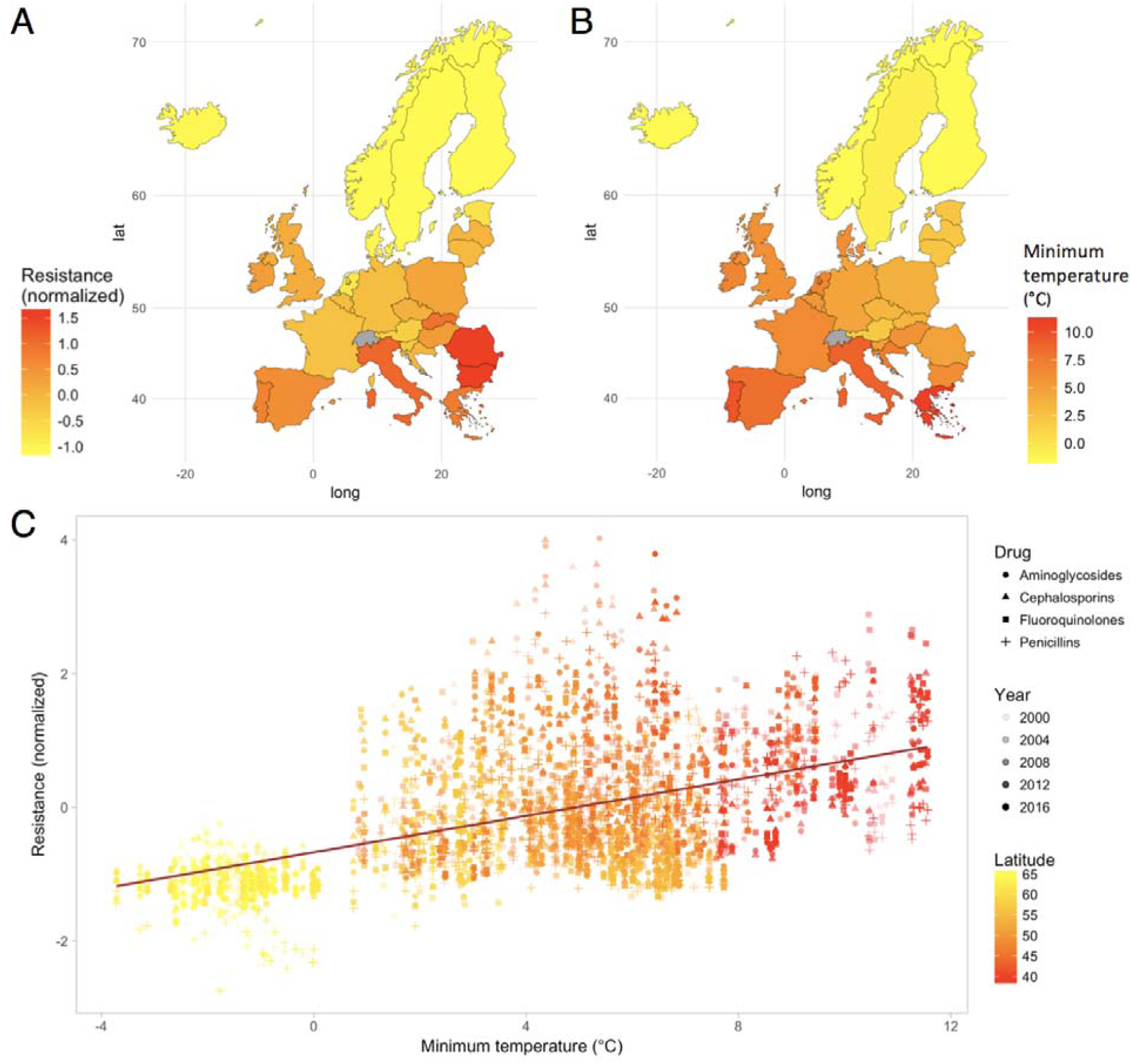
Antibiotic resistance increases with increasing minimum temperature. Maps visualizing (A) mean normalized antibiotic resistance for 3 common pathogens (*E. coli, K. pneumoniae*, and *S. aureus*) and (B) annual average minimum temperature (°C) between 2000-2016, for 28 European countries. The relationship between minimum temperature and normalized resistance, across all countries and pathogens, is shown with an unadjusted linear trend line (C). The transparency of the points in the scatter plot correspond to data collection year, and are colored by latitude. Antibiotic classes are represented as geometric shapes.

**Figure 2.**
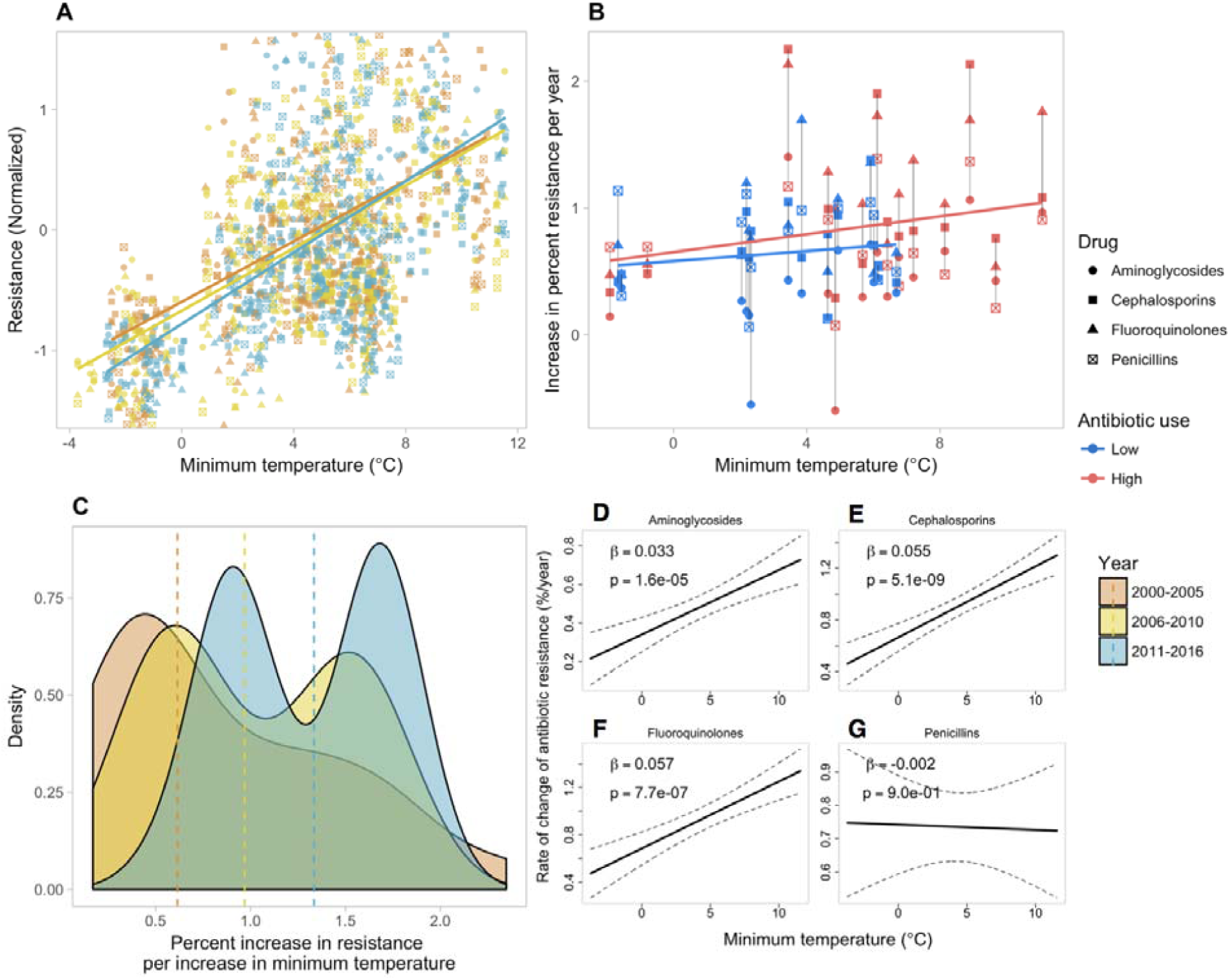
Change in the relationship between antibiotic resistance and minimum temperature over time for *E. coli*. (A) Normalized antibiotic resistance versus minimum temperature (°C) for 4 antibiotic classes and all 28 countries, stratified by 5-6 year intervals. (B) Speed of increase of antibiotic resistance over time versus average minimum temperature, stratified by average median antibiotic consumption (low/high). Each country is represented by a collection of vertical points. (C) Density distributions of unadjusted association measures (slopes) between antibiotic resistance and minimum temperature, stratified by time and with median densities (by 5-6 year intervals) marked by vertical dashed lines. (D—G) Change of antibiotic resistance over time as a function of minimum temperature for (D) aminoglycosides, (E) 3rd-generation cephalosporins, (F) fluoroquinolones, and (G) aminopenicillins, with 95% confidence intervals. Estimates for (D—G) were obtained from multivariable models adjusting for country, minimum temperature (°C), year, population density (persons/km^2^), antibiotic consumption (prescriptions/1000 inhabitants/day), and the interaction between year and minimum temperature. Beta coefficients and p-values are given for the interaction between minimum temperature and year, with 95% confidence intervals calculated using the standard error of Equation 3, where and are estimates of and.

By taking the time derivative of Equation 1, we were able to assess, for each pathogen and antibiotic class, the extent to which cross-country variations in temperature, as well as rates of change of the predictors, contributed to the rates of change of antibiotic resistance over time in different geographies. We quantified each non-intercept term in Equation 2 individually for each country and year of data, including computing the time derivatives of predictors, in order to produce a distribution of values over geographies and time. Considering these distributions, we found that differences in rates of change of antibiotic resistance across geographies were driven mainly by variations in minimum temperature, with the median of the term 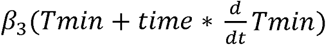 contributing at least 20 times more to the observed rates of change of antibiotic resistance, across Europe, than the terms involving temporal changes in population density, antibiotic consumption, and minimum temperature (for antibiotic classes aminoglycosides, cephalosporins, and fluoroquinolones; Fig. S9). The contribution of minimum temperature to rates of change of aminopenicillin resistance was more modest but still larger than that of the other predictors (Fig. S9). As shown in Fig. S10, the variability in minimum temperature across countries in the data is much larger than the within-country temporal variability over the 17-year time period, and countries experienced relatively small changes in population density and antibiotic consumption from year-to-year as shown in Figs. S5, S6. Thus, for each pathogen and antibiotic class, the magnitude of the term *Tmin* is substantially larger than 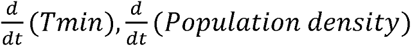 and 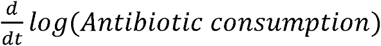 supporting why temperature is a main contributor to differences in rates of change of antibiotic resistance across Europe. Further, our analysis (Fig. S9) shows that the term *β* _3_ *Tmin* is a good estimate of the term 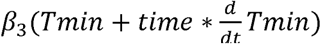, suggesting that equation 2 can be approximated by an “effective” equation given by:

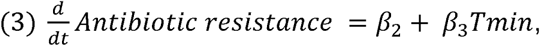

establishing the direct association between temporal changes in antibiotic resistance and local minimum temperature.

As a consequence, we found that warmer ambient minimum temperatures were generally associated with faster increases in antibiotic resistance over time, a phenomenon consistent across most pathogens and classes of antibiotics (Figs. 2D, S7D, S8D). Over the study time period and controlling for national antibiotic consumption, population density, and other constant country-specific confounders, we found that warmer countries, with 10°C larger values in average minimum temperature, experienced an increased rate of change of antibiotic resistance by 0.33%/year (p<0.001), 0.55%/year (p<0.001), and 0.57%/year (p<0.001) for *E. coli* against aminoglycosides, 3rd-generation cephalosporins, and fluoroquinolones, respectively (Table 1, Fig. 2D). We observed faster rates of change of *K. pneumoniae* resistance to 3rd-generation cephalosporins and fluoroquinolones, of 0.9%/year (p<0.01) and 1.2%/year (p<0.01) for a 10°C increase in temperature across countries, respectively (Table 1, Fig. S7D). Interestingly, we observed a decrease in the rate of change of antibiotic resistance across countries by approximately 0.4%/year (p<0.05), with a 10°C increase in minimum temperature, for the pathogen *S. aureus* and methicillin.

**Table 1:** Adjusted multivariable analyses by pathogen and antibiotic class. Coefficients with standard errors (95% confidence intervals) are adjusted for country, minimum temperature, year, population density, antibiotic consumption, and the interaction between year and minimum temperature. For interpretability, year is zeroed at baseline (2000). A log transform was applied to antibiotic consumption to improve linear fit. All available pathogen-antibiotic combinations of 3 pathogens (*E. coli*, *K. pneumoniae*, and *S. aureus*) and 4 antibiotic classes (aminoglycosides, 3rd-generation cephalosporins, fluoroquinolones, and penicillins) were analyzed. Penicillin resistance in *E.coli* was measured as resistance to aminopenicillins, and for *S. aureu* s as methicillin resistance.

|  | Aminoglycosides | Cephalosporins | Fluoroquinolones | Penicillins |
| --- | --- | --- | --- | --- |
| ***E. coli*** |  |  |  |  |
| Minimum temperature (°C) | -0.29<br>(-0.71, 0.14) | -0.22<br>(-0.74, 0.30) | -0.5<br>(-1.14, 0.13) | -0.15<br>(-0.84, 0.53) |
| Year | 0.34 ***<br>(0.25, 0.43) | 0.67 ***<br>(0.56, 0.77) | 0.68 ***<br>(0.54, 0.82) | 0.74 ***<br>(0.59, 0.89) |
| Minimum temperature (°C): Year Interaction | 0.03 ***<br>(0.02, 0.05) | 0.05 ***<br>(0.04, 0.07) | 0.06 ***<br>(0.03, 0.08) | 0<br>(-0.03, 0.02) |
| Antibiotic consumption (log DDD/1000 persons) | 0.32<br>(-0.00, 0.64) | 0.44 **<br>(0.17, 0.71) | 2.02 ***<br>(1.03, 3.00) | -0.62<br>(-2.07, 0.84) |
| Population density (persons/km<sup>2</sup>) | -0.05<br>(-0.10, 0.01) | -0.10 **<br>(-0.17, -0.03) | -0.06<br>(-0.14, 0.02) | 0.02<br>(-0.07, 0.10) |
| R<sup>2</sup> | 0.83 | 0.87 | 0.87 | 0.87 |
| ***K. pneumoniae*** |  |  |  |  |
| Minimum temperature (°C) | -1.07<br>(-2.29, 0.15) | -1.23<br>(-2.52, 0.06) | -1.34<br>(-2.93, 0.24) |  |
| Year | 0.56 **<br>(0.21, 0.91) | 0.66 ***<br>(0.29, 1.02) | 0.92 ***<br>(0.48, 1.36) |  |
| Minimum temperature (°C): Year Interaction | 0.04<br>(-0.02, 0.10) | 0.09 **<br>(0.03, 0.15) | 0.12 **<br>(0.04, 0.19) |  |
| Antibiotic consumption (log DDD/1000 persons) | 0.19<br>(-0.83, 1.20) | 0.90 **<br>(0.25, 1.54) | 3.45 **<br>(0.89, 6.01) |  |
| Population density (persons/km<sup>2</sup>) | 0.11<br>(-0.10, 0.31) | 0.04<br>(-0.17, 0.26) | -0.28 *<br>(-0.55, -0.02) |  |
| R<sup>2</sup> | 0.93 | 0.94 | 0.87 |  |
| ***S. aureus*** |  |  |  |  |
| Minimum temperature (°C) |  |  |  | 0.7<br>(-0.25, 1.65) |
| Year |  |  |  | -0.04<br>(-0.23, 0.15) |
| Minimum temperature (°C): Year Interaction |  |  |  | -0.04 *<br>(-0.07, -0.00) |
| Antibiotic consumption (log DDD/1000 persons) |  |  |  | 0.34<br>(-1.70, 2.38) |
| Population density (persons/km<sup>2</sup>) |  |  |  | -0.29 ***<br>(-0.41, -0.17) |
| R<sup>2</sup> |  |  |  | 0.87 |

\*\*\* p < 0.001; \*\* p < 0.01; \* p < 0.05.

We also observed that higher values of antibiotic consumption, thought to have important roles in conferring antibiotic resistance, including through population-level selective pressure,^10,16^ were generally associated with higher values of antibiotic resistance to cephalosporins and fluoroquinolones for the pathogens *E. coli* and *K. pneumoniae* (Table 1; Fig. S11). For a few pathogen and antibiotic combinations, lower resistance was associated with more densely populated countries (p<0.01, Table 1). Across pathogens, the relationship between antibiotic resistance and minimum temperature did not substantively change by median antibiotic consumption or by tertile of population density (Fig. S12).

In our statistical approach, we included country-specific intercepts to capture the structural differences in health and economic characteristics (including agricultural practices) across countries, which could confound the relationship between temperature and antibiotic resistance. We achieved similar effects and strong model fits with our structural model (R^2^ range: 0.81-0.94; Table 1; Table S2; Table S3) when performing the analysis using two different spatially aggregated measures of minimum temperature, namely (a) country-level, and (b) city-level, a spatial scale representative of major population activity in each country. At the city level, our findings did not substantively change when using local weather station data compared to MERRA-2 reanalysis weather data (Table S2; Table S3) or when modeling antibiotic consumption using a cubic spline (Table S4). Lastly, in order to confirm our conclusion that a long-term minimum temperature (as opposed to temporal changes of minimum temperature within countries) effect is associated with rates of change of antibiotic resistance, we removed the temporal variability in minimum temperature and replaced it with a fixed 17 -year temporal average, within each country, and repeated our analysis. As reported in Table S5, we recovered nearly identical results to the original analysis.

## Discussion

Our ecologic study presents the first evidence that the rate at which antibiotic resistance accumulates over time is associated with ambient temperature across Europe. This finding helps understand the currently observed geographic distribution of antibiotic resistance prevalence in Europe, which shows warmer regions experiencing higher levels of antibiotic resistance. We found that warmer ambient temperatures are generally associated with larger rates of increase of resistance for *E. coli* and *K. pneumoniae*, and for larger rates of decrease for *S. aureus.* These findings for *S.aureus*, which appear contradictory to those of *E. coli* and *K. pneumoniae*, may reflect concerted efforts to reduce methicillin-resistant *S. aureus* (MRSA) infections in countries with high endemic MRSA prevalence. These efforts to curb MRSA have been well documented in Europe^17^ where many countries have instituted successful programs that target healthcare-associated MRSA (HA-MRSA) transmission and infections through enhanced infection prevention and control practices. These efforts may be unique to MRSA and thus are unlikely to impact other pathogens and susceptibilities.

The role of climate and infectious diseases has been a major topic of research in the last decade. However, much of this work has focused on the impact of vector borne diseases (e.g. Malaria and Dengue Fever), as well as diseases related to water and sanitation (e.g. Cholera)^9^. To date, there has been a paucity of literature on the relationship between climate factors and population-level antibiotic resistance. An exception is an ecologic study across the United States,^10^ and due to limited temporal resolution, this study could not assess trends over time. One recent country-level analysis in Europe did not explicitly assess the effects of temperature on temporal trends in antibiotic resistance, but did find that increasing distance from the equator was a predictor of lower risk of resistance of one pathogen, *Acinetobacter spp*, to one antibiotic class, carbapenems.^18^ Compared to previous studies, our study offers evidence for the positive relationship between temperature and antibiotic resistance over a larger geographic and temporal scale, and across multiple bacterial pathogens and drug classes. Further, we focus on rates of increase to support previously observed cross-sectional relationships. Other datasets offer the future promise of evaluating the potential impact of climate on antibiotic resistance on a global scale.^5,19^

While we have a limited understanding of the transmission networks that give rise to the spread of antibiotic resistant organisms and mechanisms of resistance, there is a general understanding that this is a complex process, connecting agriculture, animals, humans, and the broader environment.^20^ Temperature plays an important role in many of these settings, and there are a number of potential mechanisms that could support a biological effect of temperature on antibiotic resistance distribution and transmission. Antibiotic resistant bacteria have either intrinsic or acquired mechanisms that render them non-susceptible to particular types of antibiotics, and these bacteria can exist in both the environment and hosts (e.g. carriage within flora of humans and animals).^21^ On a bacterial level, mechanisms of resistance can either be shared vertically (through strain replication) or horizontally (e.g. from other bacteria, viruses, or the environment).^21,22^ Temperature thus has the potential to act on any number of these resistance acquisition and transfer mechanisms, both within and between hosts and environments. For example, increased carriage in humans and animals could support increased transmission of resistant strains of bacteria, and carriage of antibiotic resistant organisms has been shown to be influenced by season/temperature.^8^ Moreover, horizontal gene transfer (including the transfer of antibiotic resistance genes between bacteria) is known to be temperature dependent, and previous work has shown that optimal temperatures for transfer of the carbapenemase NDM-1 are similar to local ambient temperatures. Increased bacterial growth rates at higher temperatures could facilitate overall transmission and selection by antibiotic consumption.^6^ The environment is thought to be a major reservoir of antibiotic resistance,^23^ and it is also possible that the environmental resistome (the collection of all resistance genes) could become more diverse in a warming climate. Lastly, temperature may have an impact on other important human socio-behavioral effects (e.g. gatherings) which could also alter the transmission of antibiotic resistant organisms.

Our results were almost identical when we conducted our statistical analysis on two different temperature data sources and two distinct spatial resolutions (country- and city-level), confirming that our findings are robust. Moreover, by removing the within-country temporal variability in minimum temperature, and conducting our analysis using a fixed 17-year average minimum temperature we recovered our original findings (Table S5). The latter result confirms that our findings establish a *long-term* effect of minimum temperature on the rate of increase of antibiotic resistance. Our attempts to identify potential short-term climatic influences of our set of predictors on antibiotic resistance did not lead to consistent patterns. For this, we conducted a post-hoc analysis on the direction of the changes, from one year to the next, of minimum temperature, antibiotic consumption, and antibiotic resistance. We calculated the proportion of times for which the direction (increase/decrease) of annual changes of antibiotic resistance and the predictors were the same. This metric, often called “hit rate,” captured the congruence in the directional movement of trends in antibiotic resistance, minimum temperature, and antibiotic consumption. Based on these short-term trends analyses, there was no general pattern suggesting that temporal deviations in temperature and antibiotic consumption, either synchronously or lagged, could explain temporal deviations in antibiotic resistance, on average across countries (mean hit rates < 60%; Fig. S13). Note that a hit rate of about 50% suggests that half of the times one of the signals increased, the other one decreased. So, a strong signal takes place when the majority of the times both move up or down. Within some countries and for some pathogens and antibiotic classes, there may be strong signatures of short-term trends (Figs. S14-S16), which future analyses may explore.

If future studies confirm that the rate at which antibiotic resistance accumulates in human pathogens is influenced by temperature at the global scale, directly or indirectly, then our findings suggest that future increases in global temperature may yield faster increases in antibiotic resistance.^24^ This finding was raised speculatively by a paper examining the geographic distribution of antibiotic resistance in the United States,^10^ and an immediate next step could involve amassing more years of data on resistance in the U.S. to confirm the findings of the present study.

Estimating the health and economic burden of antibiotic resistance globally is challenging: one extreme estimate suggests that attributable mortality due to antibiotic resistance could be on the order of ‘millions’ annually by 2050, with annual economic burden in excess of ‘billions’ of US dollars.^25–28^ Regardless of the methodology used to estimate the burden of antibiotic resistance globally, failure to account for potentially relevant factors such as temperature could lead to underestimates.

While we cannot infer causality from our study, we have demonstrated that temperature may play an important role in modulating the rate of change of antibiotic resistance in a region, and this may explain the geographic differences that have been observed in cross-sectional studies. While our approach of using country-wide minimum temperatures was done to provide comparability with national antibiotic resistance data and could lead to blunted associations, these would still likely only lead to underestimates. We hope this work will drive further avenues of research to investigate the role of climate as well as other sociodemographic factors on the distribution and transmission of antibiotic resistance.

## Acknowledgments

D.R.M. received training support from a Canadian Institutes of Health Research Fellowship grant.

## Contributors

S.F.M., D.R.M., and M.S. conceived the study; S.F.M., D.R.M., M.W.H., K.M., and M.S. formulated the experimental design; S.F.M. and D.R.M. collected the data; S.F.M., M.W.H., and M.S. analyzed the data; and all authors discussed results, contributed to manuscript preparation, and reviewed the manuscript.

## Declaration of interests

All authors declare no competing interests.

### Data and material availability

Data used in the study are from openly available sources and are described/referenced accordingly.

## Supplemental Materials

Additional Limitations

Sensitivity Analyses

Data Use Disclaimer

Figs. S1 to S16

Tables S1 to S5

